# Spatial alignment between faces and voices improves selective attention to audio-visual speech

**DOI:** 10.1101/2021.04.19.440487

**Authors:** Justin T. Fleming, Ross K. Maddox, Barbara G. Shinn-Cunningham

**Affiliations:** Department of Speech-Language-Hearing Sciences, University of Minnesota, Minneapolis, MN, U.S.A.; Speech and Hearing Bioscience and Technology Program, Harvard University, Cambridge, MA, U.S.A.; Department of Biomedical Engineering, University of Rochester, Rochester, NY, U.S.A.; Neuroscience Institute, Carnegie Mellon University, Pittsburgh, PA, U.S.A.

**Keywords:** Audio-visual, speech, attention, spatial, online experiment

## Abstract

The ability to see a talker’s face has long been known to improve speech intelligibility in noise. This perceptual benefit depends on approximate temporal alignment between the auditory and visual speech components. However, the practical role that cross-modal spatial alignment plays in integrating audio-visual (AV) speech remains unresolved, particularly when competing talkers are present. In a series of online experiments, we investigated the importance of spatial alignment between corresponding faces and voices using a paradigm that featured both acoustic masking (speech-shaped noise) and attentional demands from a competing talker. Participants selectively attended a Target Talker’s speech, then identified a word spoken by the Target Talker. In Exp. 1, we found improved task performance when the talkers’ faces were visible, but only when corresponding faces and voices were presented in the same hemifield (spatially aligned). In Exp. 2, we tested for possible influences of eye position on this result. In auditory-only conditions, directing gaze toward the distractor voice reduced performance as predicted, but this effect could not fully explain the cost of AV spatial misalignment. Finally, in Exp. 3 and 4, we show that the effect of AV spatial alignment changes with noise level, but this was limited by a floor effect: due to the use of closed-set stimuli, participants were able to perform the task relatively well using lipreading alone. However, comparison between the results of Exp. 1 and Exp. 3 suggests that the cost of AV misalignment is larger at high noise levels. Overall, these results indicate that spatial alignment between corresponding faces and voices is important for AV speech integration in attentionally demanding communication settings.

## Introduction

The ability to see a person’s face as they are speaking leads to a well-established improvement in speech recognition accuracy, particularly in high levels of background noise.^1–7^ This benefit critically depends on the temporal relationship between the auditory and visual signals. For both speech and nonspeech stimuli, temporal coherence between cross-modal features drives binding of the sensory inputs into a single perceptual object.^8^ If the signals are offset beyond the limits of a temporal binding window, benefits of audio-visual (AV) binding are abolished; for AV speech, this window spans an auditory offset of roughly −40 to 200 ms relative to the visual input.^9^ Electroencephalography studies have shown that event-related potentials (ERPs) elicited by AV speech stimuli have reduced latencies and amplitudes compared to their auditory-only counterparts, suggesting that vision can provide anticipatory information that facilitates auditory processing.^10–12^ However, as with the behavioral benefits of integration, introducing temporal offsets between the auditory and visual stimuli systematically reduces the strength of these AV ERP modulations.^13^

Consensus has not been reached regarding the practical role that AV *spatial* alignment plays in integration. On one hand, several cross-modal illusions are known to be unaffected by spatial misalignment between their unisensory components. For instance, the McGurk effect, in which visual articulator movements influence auditory perception of syllables, occurs even with a large spatial disparity between the talker’s video and voice.^14,15^ The same is true of the sound-induced flash illusion, in which the number of briefly presented auditory stimuli influences the number of perceived visual stimuli.^16,17^ However, in a study that introduced a more complex version of the sound-induced flash paradigm with multiple competing streams of auditory and visual stimuli, the strength of the effect was modulated by spatial alignment within each AV stream.^18^ Similarly, in the pip and pop effect, in which visual search is facilitated by a tone played in synchrony with a visual target,^19^ behavioral and electrophysiological signatures of integration were observed regardless of AV spatial alignment between visual targets and tones. However, in a version of the task with two competing AV stimuli, search benefits and ERP signatures of integration did depend on spatial alignment between the auditory and visual components of each stimulus.^20^ Taken together, these findings indicate that in relatively simple scenes lacking in multisensory competition, temporal coherence alone is sufficient to drive AV integration. When the sensory environment becomes more complex, however, AV spatial alignment may represent an important secondary cue to aid selective integration of the correct inputs.

Some previous studies have investigated visual facilitation of auditory selective attention using speech stimuli in “cocktail party” listening environments. For instance, visual input that is temporally coherent with one stream in an auditory mixture improves perceptual and neural tracking of that stream.^21,22^ However, to our knowledge, no existing studies have examined whether spatial alignment (or explicit misalignment) between faces and voices influences this AV selective attention advantage. In investigating this, target and distractor voices would necessarily be spatially separated from one another, providing another auditory selective attention benefit in the form of spatial release from masking (SRM).^23^ Acoustic differences in the relative levels of the target and distractor speech reaching the ears contribute to SRM, but such acoustic differences are relatively small when competing speech is present. Instead, the main benefits of spatially separating the competing talkers arise from facilitating perceptual segregation of the voices, thereby allowing listeners to focus attention selectively on the target talker based on its location.^24–26^ Given that AV binding can also improve perceptual segregation, the current work aimed to tease apart these benefits. In one previous study that combined SRM with the ability to see a target talker’s face, the presence of visual input was found to provide a greater speech recognition benefit when the target and masker speech signals were spatially coincident.^27^ However, as with most multisensory selective attention paradigms using speech stimuli, this study focused on how a *single* visual stimulus can perceptually highlight a target speech stream in an auditory mixture. The presence of competing sensory inputs in both audition and vision may provide a closer approximation of real-world communication challenges.

In the present study, we aimed to determine whether AV spatial alignment improves selective attention to speech in the presence of speech-shaped noise and a competing AV talker. Participants performed a speech selective attention task, which required them to attend a cued Target Talker while ignoring a Distractor Talker, and then indicate which of four words was spoken by the Target Talker. Importantly, multiple cues were available to separate the Target and Distractor speech streams (e.g. differences in pitch, vowel spaces, and other talker-specific characteristics), so attention to spatial features was not explicitly required to perform the task.

Data was collected online using the Gorilla online experiment platform, and sounds were spatialized with generalized HRTFs. In Exp. 1, we examined whether spatial alignment between corresponding faces and voices affected the magnitude of AV benefits in speech attention. Aligned and Misaligned AV conditions were compared against auditory-only conditions with spatially separated or spatially coincident speech streams. Exp. 2 examined gaze fixation effects on auditory spatial attention, and whether they may have influenced the results of Exp. 1. Exp. 3 tested the effect of AV spatial alignment at increasingly challenging SNRs, and Exp. 4 measured lipreading performance on this closed-set task using a visual-only condition. Across these experiments, spatial alignment between corresponding faces and voices provided a consistent selective attention benefit. While gaze position alone had an effect, it could not account for the full effect of AV spatial alignment (Exp. 2). The effect of spatial alignment between faces and voices may be magnified in noisier settings (Exp. 1 and Exp. 3), though a future open-set version of this task is needed to confirm this (Exp. 4).

## General Methods

There were several common aspects across the four experiments presented in this study. These general procedures will first be described, with experiment-specific methods covered in subsequent subsections. All experiments were presented online using the Gorilla Experiment Builder (https://www.gorilla.sc), and subject recruitment was conducted using the Prolific online recruitment service (https://www.prolific.co). Participants were required to use a laptop or desktop computer and either the Microsoft Edge or Google Chrome web browser, due to known issues with media autoplay in other web browsers. To be included in the study, participants were required to be 18-55 years old, have learned English as their first language, and have no known hearing loss. Participants were allowed to complete only one of the four experiments. Participants provided informed consent, and all study procedures were approved by the Carnegie Mellon University Institutional Review Board.

### Screening and Pre-Experiment Tasks

Before participants were allowed to start the main experiment, they had to complete several screening and setup tasks designed to ensure their web browser settings and audio equipment were suitable to perform the study. First, a questionnaire at the end of the consent form asked participants to confirm the first language and hearing status they reported in Prolific; those who failed to confirm this information were rejected. Next, a brief piece of music was automatically played to ensure that participants had autoplay enabled in their web browser. If they could not hear the music, instructions for enabling autoplay were provided. If they did not want to change their browser settings, participants were also given the option to withdraw from the study at this point.

Next, participants completed an illusory pitch detection task based on the Huggins pitch phenomenon to check that they were using headphones. In the Huggins pitch effect, identical white noise is played to the two ears, except that the noise is phase-shifted by 180° in a narrow frequency band in one ear. Monaurally, this phase shift is undetectable, but when presented dichotically, participants perceive a pitch corresponding to the narrowband noise that is phase-inverted between the ears.^28,29^ Since free-field interference disrupts this interaural phase offset, screening tasks based on Huggins’ pitch are highly selective for participants who are using headphones.^30^ On each trial, three binaural noise stimuli were presented sequentially, one of which contained a Huggins’ pitch stimulus. Participants made a 3AFC judgment about which interval contained the “hidden tone.” Participants were first given an example trial, on which they were told which interval contained the tone. They then performed six trials of the 3AFC task without feedback. All trials needed to be answered correctly to proceed to the main task, but participants were allowed one retry with six new trials if they did not pass on the first attempt.

Participants were next asked to set their computer volume in preparation for the main experiment. An audio-only speech stimulus resembling those used in the main study (two female talkers embedded in speech-shaped noise) was played, and participants were asked to turn up their computer volume until the stimulus was as loud as possible without becoming uncomfortable. Finally, participants performed a brief spatial hearing task using similar audio-only speech stimuli. Prior to starting this task, participants were instructed to check that their headphones were not on backwards. On each trial, one of the talkers from the main experiment spoke a five-word sentence, which was spatialized using generic head-related transfer functions (HRTFs) to either −15° or 15° azimuth. These sentences were not repeated by this talker in the actual experiment. Participants’ task was to judge whether the speech came from the left or the right. After two practice trials with feedback, participants were required to answer five out of six trials correct without feedback to advance to the main experiment.

### Trial Structure and Task

During the main experiment, participants were asked to pay attention to one talker (the Target Talker) while ignoring the other talker. The general timeline of a trial is illustrated in Fig. 1A. Each trial started with 1 second of fixation, with the fixation position changing based on the Target Talker and experimental condition, but remaining constant throughout the trial. Next, the word “nine” was spoken by the Target Talker to inform participants which talker to attend for the upcoming trial. The stimulus configuration was also present in the cue (e.g., video on or off, spatial location of the Target Talker’s voice). This was followed by another 1.5 s of fixation, with a separate token of speech-shaped noise (SSN) coming on in each ear for the last 500 ms. The SSN continued throughout presentation of the two competing speech stimuli. Stimulus presentation was followed by another 500 ms of fixation, after which the participant response screen.

**Figure 1:**
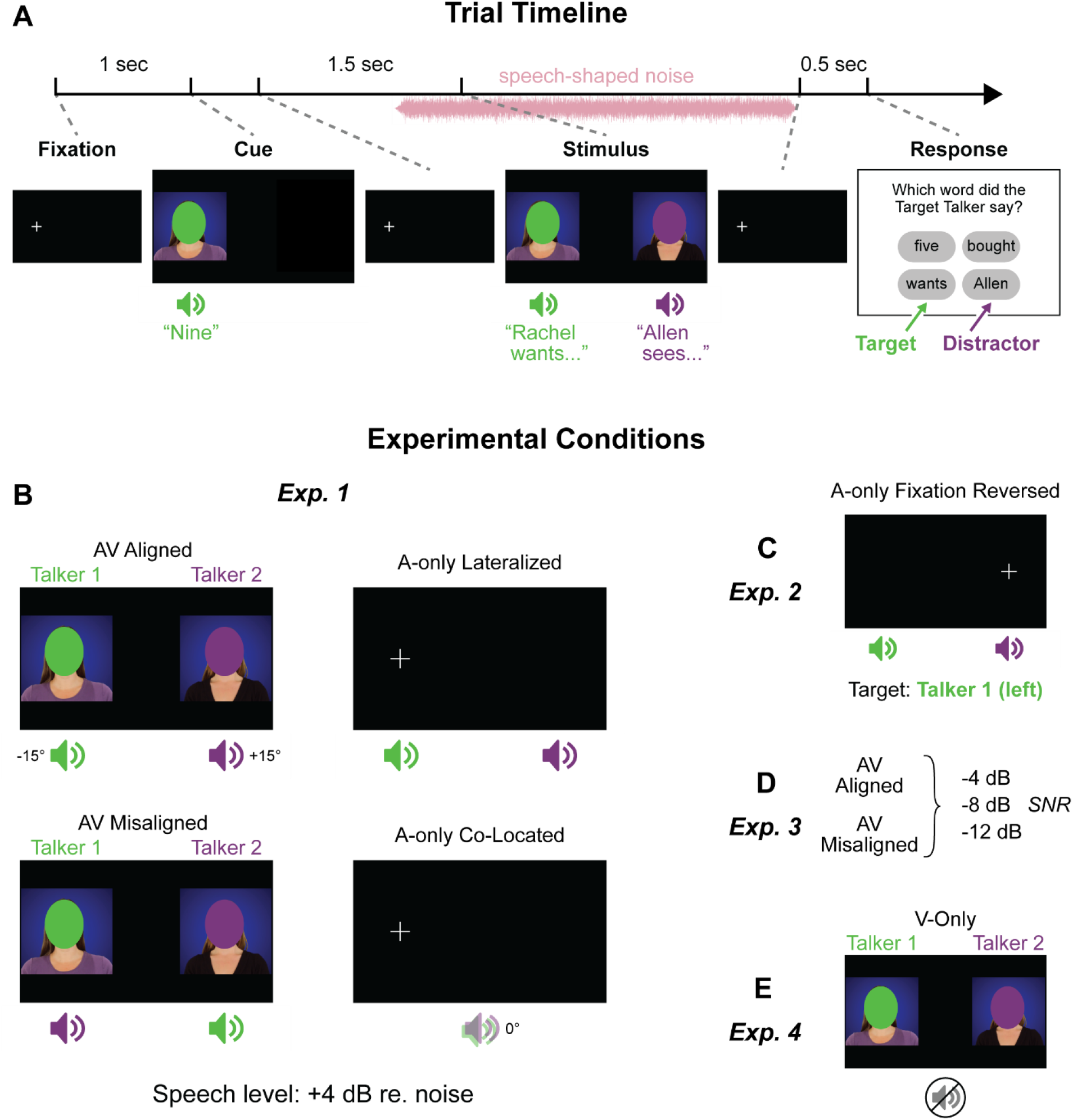
Trial timeline and illustration of experimental conditions. All panels show Talker 1 (green, left) as the Target Talker. *A*) The timeline of trial events is shown for an AV trial in which the faces and voices were spatially aligned. Audio-visual configurations for the four conditions of Exp. 1 are shown in *B*, and the new conditions added in the subsequent experiments are shown in *C-E*. Auditory stimulus locations are shown beneath the corresponding video snapshots. In Auditory-only (A-only) conditions, the fixation cross remained on the screen throughout the trial. Faces are obscured for the bioRχiv preprint.

At the response screen, participants were shown four words, one of which had been spoken by the Target Talker. Participants’ task was to identify this target word and click on it using their mouse. Among the non-target word options, one or two (chosen randomly on each trial) always came from the Distractor Talker’s stream. The other one or two word options were selected randomly from words present in the stimulus corpus, but not spoken by either talker on the current trial. The instructions stated that participants should respond as quickly as possible without sacrificing accuracy.

### Stimuli

The original speech stimuli were high-definition audio-visual recordings of two female talkers saying short sentences in a neutral affect, drawn from the Sensimetrics Speech Test Video Corpus. The video components of these stimuli are limited to the talkers’ shoulders and above and do not include hand gestures. The sentences are constrained to a syntactic structure of Name– Verb–Number–Adjective–Plural Noun (for example, “Peter gives nine red tables”). One of 10 interchangeable words can appear in each position. The corpus includes each talker saying 500 unique sentences of this structure; a subset of 160 of these sentences were chosen randomly from each talker for this experiment. On each trial, stimulus pairings from the two talkers were restricted such that all five words were different between the two talkers. Within each talker, no sentences were repeated during the experiment.

On each trial, the auditory signals from the two talkers underwent the following processing steps: amplitude normalization, onset and offset ramping, time alignment, spatialization, and the addition of speech-shaped noise. The speech envelopes were first approximated separately for each stimulus by lowpass filtering the signals (3^rd^ order Butterworth filter, 8 Hz cutoff frequency). The speech-on portions were estimated by finding the first and last points at which the envelope crossed an arbitrary threshold value. The root-mean-square (RMS) amplitude of each stimulus was calculated within the speech-on portion, and then the entire signal was scaled to an RMS level such that no clipping would occur across the entire stimulus set. This procedure ensured that the stimuli were amplitude-normalized across talkers and trials; however, given that the actual stimulus level was set by each participant in this online study, we did not control absolute sound levels.

The first and last 30 ms of the stimuli were cosine-ramped to avoid onset and offset artifacts. Within each pair of stimuli, the auditory signals were then time-aligned such that the speech-on portions overlapped maximally. Whichever signal had their earlier onset was delayed such that the midpoints of the speech-on portions matched, with the amount of time shift rounded to an integer multiple of the video frame rate to preserve AV alignment. Zeros were appended to the other signal to match stimulus lengths. The stimuli were then separately spatialized to −15°, 0°, or 15° azimuth (depending on the experimental condition) and 0° elevation by convolving them with generalized HRTFs from the CIPIC database.^31^

Speech-shaped noise was generated based on a random subset of 60 experimental stimuli (30 samples from each of the talkers). To generate the noise, we randomized and concatenated the 60 stimuli, computed the discrete Fast Fourier Transform (FFT), randomized the phases of the frequency components, and then returned the stimulus to the time domain with the inverse FFT. The resulting noise stimulus was approximately 186 s long; a random chunk of this signal was extracted for each stimulus. The level of the noise was measured by RMS, and depending on the condition, scaled to an amplitude of −4, 4, 8, or 12 dB relative to the speech signal at the louder ear. Each noise stimulus was then spatialized using the same HRTFs used previously and added to the speech stimulus.

We performed minimal additional processing on the video components of the stimuli, but to align them with the auditory stimuli, we duplicated the first or last frame until the video and audio lengths matched. Finally, the two auditory and visual (if present) stimulus components were combined in Adobe Premiere Pro. Fixation and cue trial phases were also added at this stage, and complete trials were exported as MP4 video files. To prevent lagging or freezing in a web browser-based experimental environment, the stimuli were compressed using default online video settings in the Handbrake Open Source Video Transcoder software (https://www.handbrake.fr).

### Data Analysis

Proportion correct and response time (RT) data were analyzed using mixed effects models. For the accuracy data, binomial models were employed at the level of individual trials. For the response time (RT) data, the median RT was first calculated for each participant and condition to limit the influence of anomalously slow RTs. These median RTs were then analyzed using linear mixed effects models. In both cases, the significance of fixed effects terms and their interactions (where appropriate) were assessed by subjecting the model to a Type III ANOVA, with p-values based on the Satterthwaite approximation for degrees of freedom. Post-hoc comparisons between each relevant pair of factor levels were made by extracting model terms for these contrasts, then cycling which level was treatment-coded as baseline until all the necessary contrasts were computed. To account for multiple comparisons, the p-value criterion for assessing the significance of these model terms was adjusted using the Bonferroni-Holm correction, assuming a starting alpha level of p = 0.05.

For plotting, each participant’s median RT data were centered on the average of their median RTs across conditions due to the large inter-subject variability in RT. Stimulus processing and data organization were performed in Python, while statistical analyses (using the lme4 and lmerTest packages) and figure generation were conducted in R.

## Experiment 1 Methods

### Participants

100 participants completed all experimental procedures for Exp. 1. 10 additional participants completed some but not all components; two failed to confirm either their first language or hearing status as reported in Prolific, six failed one of the headphone checks and were rejected, and two started the main experiment but did not finish it. Of the 100 complete datasets, four were rejected because participants performed below chance in one or more of the experimental conditions, making it difficult to verify that they were consistently attending the correct talker. Thus, the final dataset comprised 96 participants (mean age = 30.1 years, SD = 8.8; 57 female, 38 male, 1 non-binary). Participants who failed headphone screenings or did not complete the task due to technical issues were awarded partial compensation. Participants who completed all study components were paid a flat amount of $5.50, corresponding to an average pay rate of $7.77/hour. *Experiment Design*

Participants performed the speech selective attention task in four different conditions (Fig. 1B). In the *AV Aligned* condition (top-left panel), participants heard two competing speech streams and saw corresponding video of the talkers’ faces. The videos were positioned on opposite sides of the screen, with each talker’s video appearing in the same location on all trials. The auditory signals were spatialized to −15° and 15° such that each voice was presented in the same hemifield as the corresponding face. In the *AV Misaligned* condition (bottom-left panel), the video components were structured identically to the AV Aligned condition (with the same talker’s face always appearing on the left), but the voice spatialization was reversed, such that each voice was presented in the opposite hemifield as the corresponding face. In both AV conditions, participants were instructed to look at the Target Talker’s face in whatever way felt natural.

Two auditory-only (A-only) control conditions were also included so we could test for perceptual advantages of being able to see the talkers’ faces. In the *A-only Lateralized* condition (Fig. 1B, top-right panel), voices were spatialized in the same was as in the AV Aligned condition, but the video components were removed and replaced with a constant fixation cross. Fixation was lateralized to the same hemifield as the target voice to control for possible eye position effects on auditory localization and attention.^32,33^ Finally, in the *A-only Co-Located* condition (bottom-right panel), both voices were spatialized to 0° azimuth, which removed the benefit of SRM.^23^ In this condition, the fixation cross was presented in the “stereotyped” hemifield for the Target Talker (i.e., where their voice was presented in the AV Aligned and A-only Lateralized conditions) such that fixation would be lateralized similarly across conditions.

In all experimental conditions, a separate token of speech shaped noise drawn for each talker, spatialized to the same location as that talker’s voice, and set to be 4 dB quieter than the speech. Prior to starting the main experiment, participants were given one practice trial from each condition with feedback. If they answered a practice trial incorrectly, they were asked to repeat the trial until they correctly chose the target word. Trials from the four conditions were randomly intermixed throughout the experiment. Participants performed 40 trials of each condition (160 trials total) without feedback. An opportunity to take a break was provided every 10 trials.

## Experiment 1 Results

### Speech Attention Task Accuracy

A binomial mixed effects model was used to analyze task accuracy. The model had one fixed effect term, Condition, with the four experimental conditions as levels. Random effects included participant-specific intercepts for each condition and a Stimulus term. A fuller version of the model also included a random effect term for participant age, but removing this term did not result in a significantly worse fit to the data as measured by AIC and BIC, and so the simpler model was favored. The structure of the final model was as follows, in Wilkinson notation:

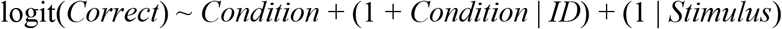

An ANOVA conducted on this model revealed a strong main effect of Condition (Χ^2^ = 204.58, df = 3, p = 4.33 · 10^−44^; Fig. 2A). Post-hoc comparisons (Wald’s z tests on model contrasts, followed by Bonferroni-Holm correction of the alpha criterion) revealed that participants performed significantly worse on the A-only Co-Located condition than any of the other three (vs. AV Aligned, z = −13.70, p = 1.03 · 10^−42^; vs. AV Misaligned, z = −8.96, p = 3.02 · 10^−19^; vs. A-only Lateralized, z = −9.60, p = 7.78 · 10^−22^). The difference between the two A-only conditions validates a strong benefit of SRM, on the order of a 30% improvement in target speech recognition, using an online platform. To assess AV benefits beyond spatial separation of the two voices, each AV condition was compared against the A-only Lateralized condition. We observed an additional performance benefit of seeing the talkers’ faces when each voice was presented in the same hemifield as the corresponding face (AV Aligned vs. A-only Lateralized, z = 4.72, p = 2.39 · 10^−6^). On the other hand, speech recognition accuracy did not differ significantly between the AV Misaligned and A-only Lateralized conditions (z = −0.40, p = 0.68). A direct comparison between the two AV conditions revealed that accuracy was significantly higher when the unisensory components of AV speech were spatially aligned than when they were misaligned (AV Aligned vs AV Misaligned, z = 4.87, p = 1.09 · 10^−6^). All significant p-values survived Bonferroni-Holm adjustment of the alpha criterion for significance.

**Figure 2:**
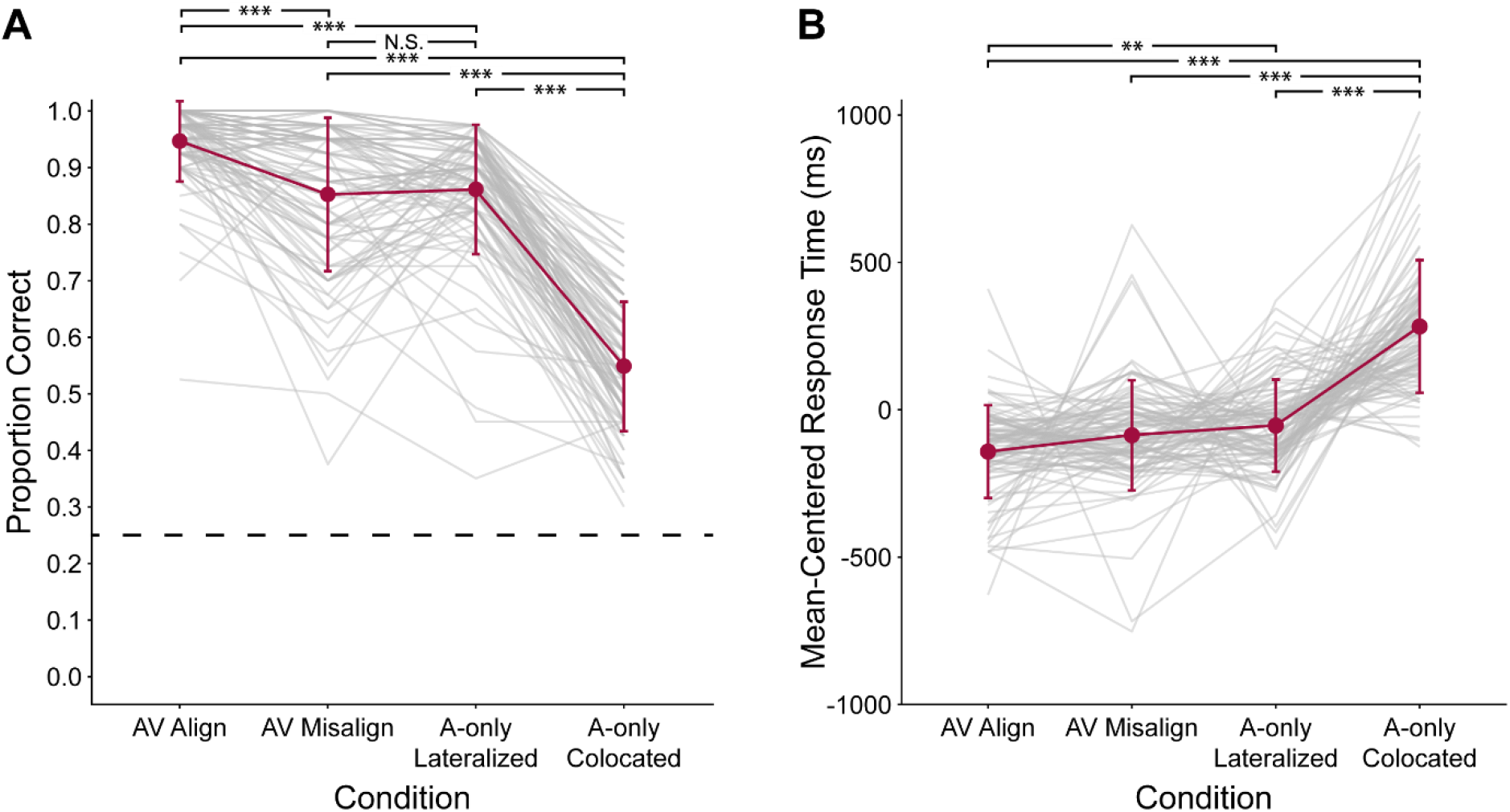
Task performance in Exp. 1. *A*) Proportion correct. Chance performance is indicated by the dashed line. *B*) Response time, with each participant’s average RT across conditions subtracted from their RT in each condition. Grey lines represent individual participants, error bars represent standard deviation, and stars indicate statistical significance in post-hoc comparisons: ** = p < 0.01, *** = p < 0.001, N.S. = not significant.

### Response Time

Median RTs within each condition were widely variable across participants, from a minimum value of 1224 ms to a maximum of 3911 ms. Nonetheless, the pattern of RTs varied systematically across conditions in a matter consistent with the accuracy data. Since the RT data were first reduced to the median across stimuli within each condition, these data were modelled using a linear mixed effects model with no random effects term for stimulus. The structure of this model was:

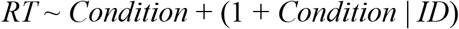

An ANOVA conducted on this model again revealed a significant main effect of Condition (Χ^2^ = 235.73, df = 3, p = 2.97 · 10^−51^; Fig. 2B). Post-hoc testing showed that participants were slower to respond in the A-only Co-Located condition than any of the other three, reflecting elevated task difficulty when neither spatial separation nor visual information were available to help segregate the speech streams (vs. A-only Lateralized, t = 10.99, p = 1.17 · 10^−23^; vs. AV Misaligned, t = 12.06, p = 2.49 · 10^−27^; vs. AV Aligned, t = 13.88, p = 8.47 · 10^−34^). A modest difference in RT was also observed between the AV Aligned and A-only Lateralized conditions (mean RTs of 1999 ms and 2087 ms, respectively), which was statistically significant (z = −2.88, p = 0.0042, Bonferroni-Holm adjusted p-value criterion: p = 0.017). This AV facilitation of RT did not reach significance when comparing the AV Misaligned and A-only Lateralized conditions (t = 1.07, p = 0.29), hinting at a particular AV RT benefit when faces and voices were spatially aligned. However, the RT difference between the AV Aligned and AV Misaligned conditions approached but did not reach significance in this experiment (t = 1.82, p = 0.07, adjusted p-value criterion: p = 0.025).

## Experiment 2 Methods

In the previous experiment, performance differences between conditions may have been partially caused by eye position, which has been shown to affect auditory localization^34,35^ and auditory selective attention.^32^ In the AV Aligned and A-only Lateralized conditions, participants gaze was held in the same hemifield to which they were listening. On the other hand, in the A-only Co-Located condition, participants fixated laterally but listened to a target voice presented at the midline, and in the AV Misaligned condition, participants looked at the Target Talker’s face in the opposite hemifield as the corresponding voice. Directing gaze toward an auditory distractor stream in this manner can reduce participants’ ability to selectively attend and remember information in an auditory target stream.^33^

To account for these issues, we conducted a follow-up experiment in which the A-only Co-Located condition was replaced with an A-only Fixation Reversed condition. On these trials, target and distractor voices were spatialized as in the A-only Lateralized condition, but participants were asked to fixate in the opposite hemifield as the target voice. Poorer performance in this condition than the A-only Lateralized condition would provide evidence that eye position effects may have contributed the difference between the AV Aligned and AV Misaligned conditions observed in Exp. 1. Much of the methodology was the same as in Exp. 1, and so this section will focus on differences between the two experiments.

### Participants

109 participants were included in the final dataset for Exp. 2 (mean age = 30.0 years, SD = 8.8; 58 female, 49 male, 2 non-binary), none of whom had participated in Exp. 1. A total of 52 additional participants were rejected for the following reasons: four failed to verify their language and hearing information from Prolific, 17 failed the headphone screening task, eight started but did not complete the main task, seven were rejected due to below-chance performance in one or more experimental conditions (6 in the new A-only Fixation Reversed condition, 1 in the AV Misaligned condition), and 16 failed a new fixation check sub-task implemented in this experiment (more below). Including these rejected participants, 161 individuals were originally recruited into the experiment. Below-chance performers were paid in full, and the remaining rejected participants received partial compensation, maxing out at $3 for those who completed all study procedures but failed the fixation check. Full payment was a flat $5.50, yielding an average pay rate of $7.53 per hour.

### Experiment Design

The AV Aligned, AV Misaligned, and A-only Lateralized conditions from Exp. 1 were left largely unchanged in Exp. 2. In the new A-only Fixation Reversed condition, the talkers’ voices were again spatialized horizontally to −15° and 15°, but the fixation cross was positioned in the hemifield opposite the target talker’s voice (see Fig. 1C). In all four conditions, catch trials were included to ensure that participants were maintaining fixation in the correct location. On these trials, a small green dot was briefly presented at the center of the fixation cross (A-only conditions) or the target talker’s video (AV conditions). The dot was presented for 300 ms at a random time when both talkers were speaking. When participants saw this dot, they were instructed to ignore the normal speech attention task and click a separate “catch” button on the response screen. Four catch trials replaced actual trials in each condition (16 catch trials overall), so participants performed 36 actual trials of each experimental condition. Participants were required to detect at least 60% of the catch trials overall, and at least 2 out of the 4 in each condition, to be included in the dataset and receive full payment.

## Experiment 2 Results

### Speech Attention Task Accuracy

A binomial mixed effects model was again used to analyze task accuracy. The conditions in this experiment were separated into two factors: Modality (AV or A-Only) and Fixation (Aligned or Reversed). Note that in the AV conditions, this Fixation term also captured effects of AV spatial alignment. The model included these factors and their interaction as fixed effects, and random effect terms for participant-specific intercepts and individual stimuli:

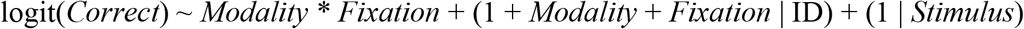

Since this experiment had a two-by-two factorial design, the model was first computed with sum-coded contrasts, such that the significance of fixed effect model terms could be interpreted in a similar fashion to ANOVA results. The Modality term was significant (z = 4.96, p = 7.22 · 10^−7^), reflecting better performance in the AV than the A-only conditions. The Fixation term was also significant (z = 6.69, p = 2.24 · 10^−11^), indicating that participants generally performed better when fixating in the same hemifield as they were listening (Fig. 3A).

**Figure 3:**
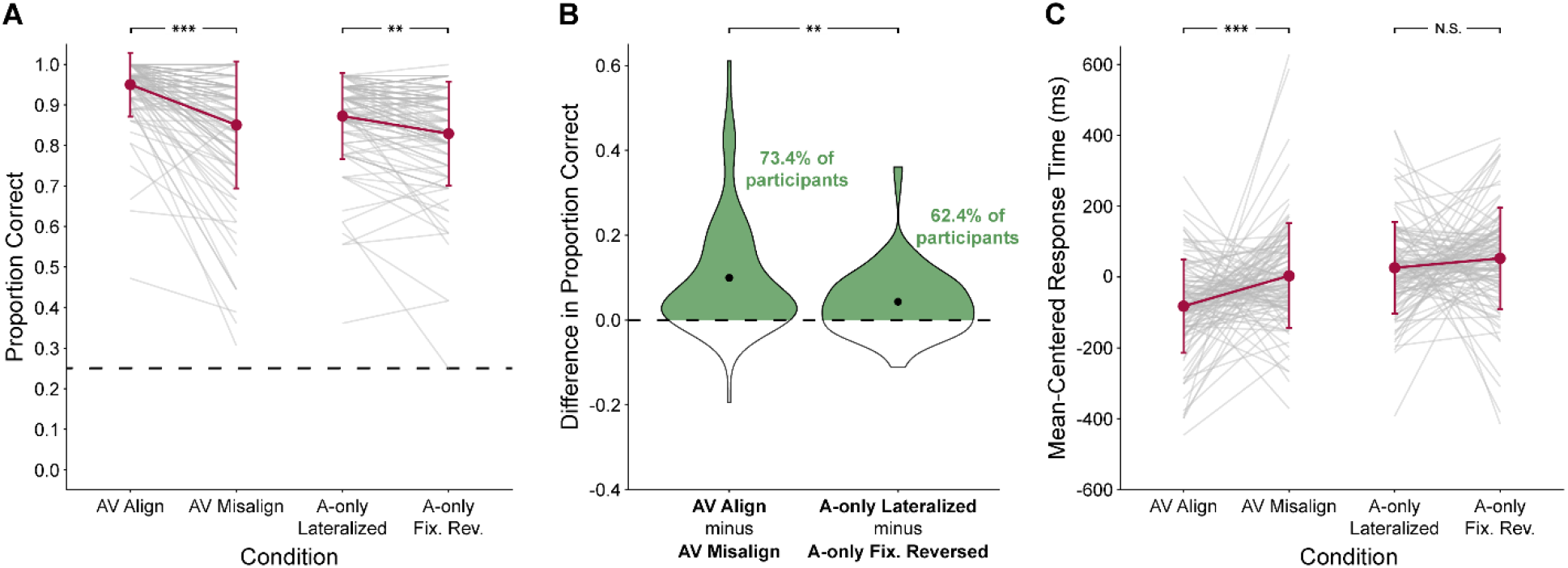
Task performance in Exp. 2. *A*) Proportion correct. Chance performance is indicated by the dashed line. *B*) To illustrate the interaction between the Modality and Fixation terms, the difference in proportion correct between the two Fixation levels is shown for the AV and A-only conditions. Black dots represent means of the distributions. *C*) Response time, with each participant’s average RT across conditions subtracted from their RT in each condition. Grey lines represent individual participants, error bars represent standard deviation, and stars indicate statistical significance in post-hoc comparisons: ** = p < 0.01, *** = p < 0.001, N.S. = not significant.

Importantly, the interaction term between Modality and Fixation was also significant (z = 2.63, p = 0.009), indicating that the effect of spatial alignment between faces and voices in the AV conditions was larger than the gaze position effect in the A-only conditions (Fig. 3B). However, post-hoc testing revealed significant effects of gaze position in both the AV (AV Aligned vs. AV Misaligned, z = 6.51, p = 7.62 · 10^−11^) and A-only (A-only Lateralized vs. A-only Fixation Reversed, z = 3.07, p = 0.002) conditions. Thus, eye position effects may have contributed to the performance difference between the AV Aligned and AV Misaligned conditions, but the interaction term indicates an added detrimental effect of having to attend auditory and visual speech signals across hemifields.

### Response Time

RTs were modeled using a linear mixed effects model with Modality, Fixation, and their interaction included as the fixed effect terms:

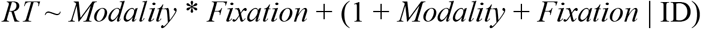

This model revealed that RTs were significantly impacted by Modality (t = −5.14, p = 1.27 ·10^−6^), with faster RTs in the AV than the A-Only conditions, and Fixation (t = −3.49, p = 6.95 · 10^−4^), with generally faster RTs when fixation was aligned with the target voice than when it was misaligned, both in accord with the accuracy data (Fig. 3C). The interaction term also reached marginal significance (t = 2.02, p = 0.046), motivating post-hoc tests. These tests revealed that RTs were significantly faster in the AV Aligned condition than the AV Misaligned condition (t = −3.94, p = 1.09 · 10^−4^), but that fixation did not significantly affect RTs in the A-only conditions (t = −1.22, p = 0.22). Thus, RTs in this speech selective attention task were speeded by the ability to see the talkers’ faces, particularly when the cross-modal components of the target and distractor streams were spatially aligned. The significant RT difference in the AV conditions, but not the A- only conditions, suggests an attentional cost when audio-visual speech is separated across hemifields, which cannot be fully explained by gaze differences.

## Experiment 3 Methods

In Exp. 1 and 2, near-ceiling performance was observed in the AV Aligned condition. Significant effects of audio-visual spatial alignment were found in both experiments in spite of this ceiling effect, but we reasoned that more dramatic effects of alignment between faces and voices would be observed if the task were made more difficult. Such an effect could also be interpreted as an example of inverse effectiveness, a principle of multisensory integration which states that the greatest multisensory gain is observed when the unisensory stimulus components elicit weak behavioral or neuronal responses.^3,36–38^ As louder noise corrupts the envelope of the speech signals, which likely contains the temporal information that binds each voice to the corresponding face, we expected that AV spatial alignment would take on greater importance as a secondary cue to AV speech integration. To this end, in Exp. 3 we tested effects of AV spatial alignment at varying SNRs, all of which were more difficult than those used in Exp. 1 and 2.

### Participants

100 participants were included in the final dataset for Exp. 3 (mean age = 31.8 years, SD = 10.2; 52 female, 47 male, 1 non-binary), none of whom had participated in the previous experiments. A total of 26 additional participants were rejected for the following reasons: nine failed the headphone screening task, four started but did not complete the main task, and 13 failed the fixation check sub-task. Since more difficult conditions were being introduced, we no longer excluded data on the basis of below-chance performance in any of the experimental conditions. As with the previous experiments, rejected participants received partial compensation and the full payment amount was $5.50, yielding an average pay rate in this experiment of $8.37 per hour.

### Experiment Design

This experiment used only the AV Aligned and AV Misaligned conditions. Whereas the noise level was previously set to be 4 dB quieter than the speech, in this experiment we tested three more difficult SNRs of −4, −8, and −12 dB. The different SNRs were achieved by varying the noise level while keeping the stimulus level constant. Prior to the experiment, an extra trial from the −12 dB condition (the loudest condition overall) was used to let participants set their system volume. Fixation catch trials, as introduced in Exp. 2, were also used here. Nine catch trials were included in both the AV Aligned and AV Misaligned conditions (18 overall, evenly distributed across SNRs). Only sessions in which participants got six out of nine catch trials correct in each AV spatial alignment condition were included in the final dataset. In addition to the catch trials, participants performed 23 trials of each combination of AV spatial alignment and SNR, making for a total of 156 trials. Conditions were randomly intermixed throughout the experiment.

## Experiment 3 Results

### Speech Attention Task Accuracy

A binomial mixed effects model was used to analyze task accuracy. The conditions in this experiment were separated into two factors: AV Spatial Alignment (Aligned or Misaligned) and SNR (−4, −8, or −12). The model included these factors and their interaction as fixed effects, and random effect terms for participant-specific intercepts and individual stimuli:

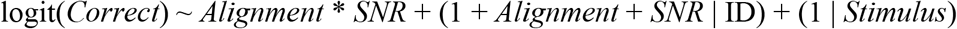

Significance of these fixed effects was assessed by passing the model to an ANOVA, with p-values based on the Satterthwaite approximation for degrees of freedom. This ANOVA revealed significant main effects of AV Spatial Alignment (Χ^2^ = 15.41, df = 1, p = 8.65 · 10^−5^), with better performance in the AV Aligned condition, and SNR (Χ^2^ = 10.95, df = 2, p = 0.004), with performance worsening as the SNR was lowered. There was also a marginally significant interaction between AV Spatial Alignment and SNR (Χ^2^ = 5.97, df = 2, p = 0.051; Fig. 4A).

**Figure 4:**
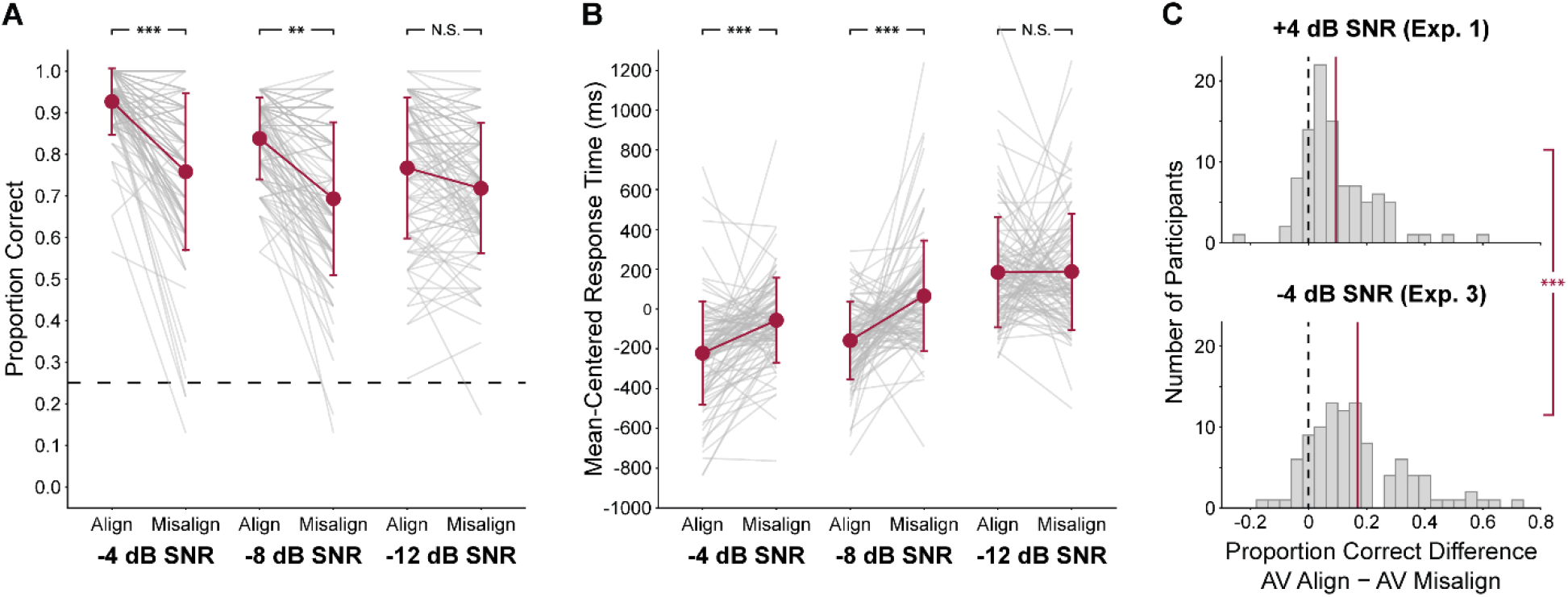
Task performance in Exp. 3 and comparison to Exp. 1. *A*) Proportion correct. Chance performance is indicated by the dashed line. *B*) Response time, with each participant’s average RT across conditions subtracted from their RT in each condition. Grey lines represent individual participants, error bars represent standard deviation, and stars indicate statistical significance in post-hoc comparisons: ** = p < 0.01, *** = p < 0.001, N.S. = not significant. *C*) Histograms of the difference in proportion correct between the AV Aligned and AV Misaligned conditions at +4 dB SNR (data from Exp. 1) and −4 dB SNR. The vertical dashed lines indicate no difference between these conditions, while each red line indicates the average difference across the participant population. Stars indicate significance in a Wilcoxon rank-sum test: *** = p < 0.001.

Post-hoc testing was restricted to comparisons between the AV Aligned and AV Misaligned conditions at each SNR, as we were interested in how this effect changed as a function of SNR. Performance on the speech attention task was significantly better in the AV Aligned than the AV Misaligned condition at −4 dB SNR (z = 3.84, p = 1.24 · 10^−4^) and −8 dB SNR (z = 2.62, p = 0.009), but not at the most difficult SNR, −12 dB (z = 0.51, p = 0.61). This pattern ran counter to our hypothesis, with effects of AV spatial alignment appearing to weaken as the task was made more difficult; a likely explanation for this will be discussed below.

### Response Time

RTs were modeled using a linear mixed effects model with AV Spatial Alignment, SNR, and their interaction included as fixed effect terms:

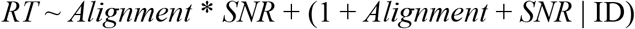

Similar to the accuracy data, submitting this model to an ANOVA revealed main effects of AV Spatial Alignment (Χ^2^ = 27.57, df = 1, p = 1.51 · 10^−7^), SNR (Χ^2^ = 87.26, df = 2, p = 2.2 · 10^−16^), and an interaction between these factors (Χ^2^ = 24.80, df = 2, p = 4.11 · 10^−6^; Fig. 4B). Post-hoc testing revealed significantly faster response times in the AV Aligned than the AV Misaligned condition at −4 dB SNR (t = −4.54, p = 8.02 · 10^−6^) and −8 dB SNR (t = −6.16, p = 2.41 · 10^−9^), both of which survived Bonferroni-Holm adjustment of the significance criterion, but not at −12 dB SNR (t = −0.09, p = 0.93). Thus, the RT results mirrored the accuracy data, with AV spatial alignment significantly speeding responses at the relatively high SNRs, but not at the lowest SNR, when responses were slowest overall.

### Comparison of SNRs across Exp. 1 and Exp. 3

Participants appeared to hit the performance floor at a much higher rate than chance, which will be examined directly in Exp. 4. This likely artificially limited effects of AV spatial alignment at the lowest SNR, as we were unable to push performance below around 70% correct. However, at +4 dB SNR (Exp. 1) and −4 dB SNR (Exp. 3), average performance in both alignment conditions was above this floor level of performance. Thus, we next compared the effect of AV spatial alignment between these two SNR levels across the experiments (Fig. 4C). At both SNRs, proportion correct was computed for each participant in the AV Aligned and AV Misaligned conditions. These proportion correct scores were then subtracted, yielding the individual benefit (in terms of a proportion correct difference) of spatial alignment between corresponding faces and voices. Shapiro-Wilk tests revealed that these difference scores were not normally distributed across the population at either SNR; thus, the effect of AV spatial alignment was compared between the +4 dB and −4 dB SNR conditions using a two-sided Wilcoxon rank-sum test. This test showed a greater benefit of AV spatial alignment at the lower SNR from Exp. 3 (W = 3480, p = 8.74 · 10^−4^). This result should be interpreted carefully, as there was a potential ceiling effect in the AV Aligned condition in Exp. 1, although average performance in the AV Aligned condition was similar between these two SNRs (94.7% correct at +4 dB SNR, 92.7% correct at −4 dB SNR). This provides some evidence that effects of AV spatial alignment may indeed become stronger as the auditory speech signal is degraded, but this should be validated using a paradigm in which proportion correct scores can be reduced to a greater extent.

## Experiment 4 Methods

The floor effect observed in Exp. 3 could have resulted from the closed-set nature of the stimuli and response method used in the current study. Although neither talker ever repeated the same sentence during the experiment, the words in the corpus were all repeated, opening the possibility that participants learned the set of possible words during the experiment. Since participants had to select the target word rather than repeating what they heard, they essentially received feedback on this learning at the response stage. While introducing a different stimulus set or response methodology were beyond the scope of the current study, in Exp. 4 we aimed to assess the extent to which these closed-set factors influenced our results. One way that this could manifest is in an increased ability to lipread the stimuli given prior knowledge of the possible words in the corpus. If this were the case, participants may have been encouraged to rely more on lipreading at lower SNRs, rendering spatial alignment between the faces and voices irrelevant. Here, we performed a version of Exp. 3 with the addition of a visual-only (V-only) condition to determine the performance level participants were able to achieve using lipreading alone.

### Participants

39 participants were included in the dataset for Exp. 4 (mean age = 34.7 years, SD = 10.1; 15 female, 24 male, 0 non-binary), none of whom had participated in the previous experiments. A total of 17 additional participants were rejected for the following reasons: two failed to confirm their native language information from Prolific, 11 failed the headphone screening task, two started but did not complete the main task, and two failed the fixation check sub-task. Data was not excluded on the basis of below-chance performance. As with the previous experiments, rejected participants received partial compensation and the full payment amount was $5.50, yielding an average pay rate in this experiment of $8.56 per hour.

### Experiment Design

The AV Aligned and AV Misaligned conditions were again used, keeping the −4 and −12 dB SNR conditions from Exp. 3. In place of the −8 dB SNR condition, we added V-only trials on which participants attempted to lipread the Target Talker’s speech with no voice. Fixation catch trials were included as in Exp. 2 and 3. Three catch trials were included in each combination of AV spatial alignment and SNR, as well as the V-only condition (15 catch trials overall). Only sessions in which participants got two out of three catch trials correct in each condition were included in the final dataset. In addition to the catch trials, participants performed 23 trials of each condition, making for a total of 130 trials. To maximize potential learning of the words in the corpus, participants first performed intermixed AV Aligned and AV Misaligned trials in the −4 dB SNR condition. We reasoned that participants would be most likely to hear all the words spoken by the Target Talker – and possibly some spoken by the Distractor Talker – at this relatively high SNR. Participants then completed the remaining AV trials in the −8 and −12 dB SNR conditions, as well as the V-only trials, all randomly intermixed.

## Experiment 4 Results

### Speech Attention Task Accuracy

Performance was quite high on average in the V-only condition (66.8% correct, SD = 18.4%). Previous studies have shown that, in normal-hearing participants, open-set word recognition performance using lipreading is typically between 10 and 15% correct.^4,39^ A binomial mixed effects model was used to analyze differences between experimental conditions, which were all coded as a single Condition factor. The model included Condition as the only fixed effect, and random effect terms for participant-specific intercepts and individual stimuli:

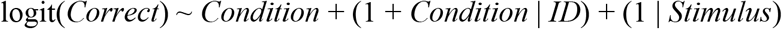

Significance of the Condition effect was assessed by passing the model to an ANOVA, with p-values based on the Satterthwaite approximation for degrees of freedom. A significant effect of Condition was found (Χ^2^ = 20.67, df = 4, p = 3.68 · 10^−4^, Fig. 5). Post-hoc tests compared the V-only condition to each of the other four, as well as the AV Aligned vs. AV Misaligned conditions within each SNR level (six total comparisons). The latter comparisons replicated the results of Exp. 3, with performance significantly improved when faces and voices were spatially aligned at −4 dB SNR (z = 2.97, p = 0.003; adjusted p-value criterion = 0.01), but not at −12 dB SNR (z = 0.75, p = 0.45). When the auditory components of AV speech stimuli were presented at −12 dB SNR, task performance was not significantly different than when participants used only lipreading, regardless of AV spatial alignment. The same was true of the AV Misaligned speech at −4 dB SNR; only AV Aligned task performance at this SNR was significantly better than the V-only condition (z = 4.36, p = 1.32 · 10^−5^; adjusted p-value criterion = 0.008). In sum, this experiment revealed that participants could achieve a high-level of lipreading accuracy, setting a floor level of performance that participants approached when the auditory stimuli were embedded in noise at −12 dB SNR.

**Figure 5:**
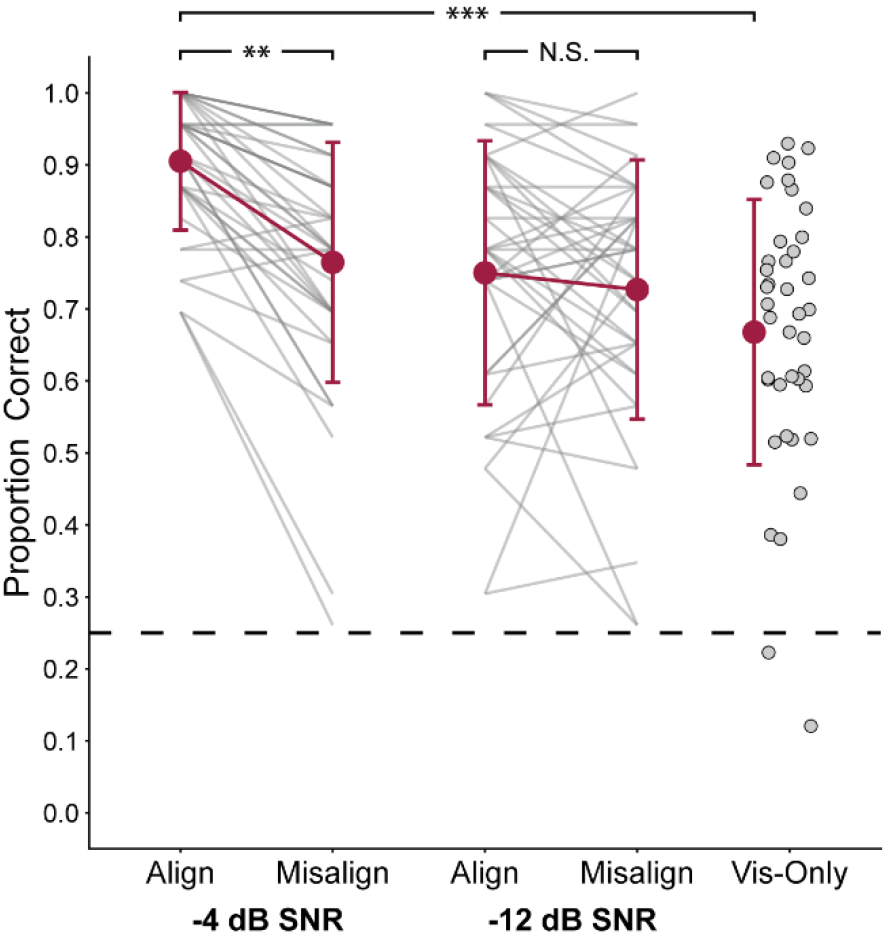
Task performance in Exp. 4. Proportion correct scores are shown, with chance performance indicated by the dashed line. Grey lines and dots represent individual participants, error bars represent standard deviation, and stars indicate statistical significance in post-hoc comparisons: ** = p < 0.01, *** = p < 0.001, N.S. = not significant.

## General Discussion

Across four online experiments, we evaluated the extent to which AV speech perception benefits depend on spatial alignment between faces and voices, using a paradigm that featured both acoustic noise and a competing talker. In Exp. 1, performance was worst overall by far in the A-only Co-Located condition, in which neither visual information nor spatial separation of the talkers were available to help participants segregate the speech streams. This indicates that the benefits of SRM in a multi-talker environment were preserved in the online experiment format.^40–42^ Beyond this benefit of spatial separation, the ability to see the talkers’ faces further improved task performance, but critically, this was only true when the corresponding faces and voices for each talker were aligned in the same hemifield. In Exp. 2, we examined the possibility that fixating in one hemifield while listening to a target in the other led to the performance decrement in the AV Misaligned condition. Indeed, auditory-only task performance was worse when participants fixed their gaze in the hemifield of the Distractor Talker, but this effect was smaller than the effect of spatial alignment between faces and voices in the AV conditions. The AV spatial alignment effect decreased as the SNR was reduced from −4 to −12 dB (Exp. 3), but participants may have hit the performance floor at the lowest SNR, as lipreading performance was quite good given the closed-set nature of the stimuli (Exp. 4). Nonetheless, comparison of the alignment effect in the +4 dB SNR condition of Exp. 1 and the −4 dB condition of Exp. 3 revealed a stronger effect at the lower SNR. This suggests that AV spatial alignment may become a more relevant cue as noise degrades the speech envelope, which provides temporal information linking the voice to the corresponding face.

Despite quite large inter-subject differences in overall response times, consistent patterns emerged across conditions in line with the performance accuracy results. The large degree of inter-subject differences was likely caused by a combination of variability in participants’ personal hardware and the response method of clicking on the word spoken by the target talker; the starting position of the participant’s mouse and the order in which they examined the possible words both could introduce noise into RT measures. In the first two experiments, RTs were reliably faster in the AV than the A-only conditions, in general agreement with task accuracy. Faster responses to multisensory as opposed to unisensory stimulation is a hallmark of multisensory integration, at least when simple stimuli are used.^43^ However, when more temporally complex AV stimuli (including speech) are used, it has been reported that AV RTs are actually slower than those to A-only stimuli.^44,45^ Thus, the speeded AV responses we observed probably do not reflect differences in the speed of early sensory processing; a more likely explanation is that AV integration made the task easier by facilitating the allocation of selective attention to the target talker.

### Spatial attention in audio-visual processing

A broad literature has converged to demonstrate that top-down attention influences the strength of multisensory integration.^46^ This has been frequently shown in the realm of AV speech using McGurk effect manipulations.^47^ If, instead of the standard single face, two lateralized faces accompany a single auditory stimulus, the face to which covert attention is directed has greater influence over auditory syllable perception than the unattended face.^48^ Similar results were found when the long-term focus of spatial attention was shifted by reliably presenting auditory stimuli from a particular location (e.g., presenting stimuli from −90° azimuth on 90% of trials); the McGurk percept was strengthened at this attended location.^49^ Neurally, deploying top-down spatial attention to AV stimuli modulates ERPs elicited by them in several time ranges, indicating effects at multiple processing stages.^50^ Similarly, fMRI activation differs across a wide network depending on whether visuo-spatial attention is directed toward speaking lips that are matched or unmatched to an auditory speech stimulus.^51^ These studies, among others, demonstrate an unequivocal link between whether a cross-modal stimulus is attended and whether signatures of multisensory integration are observed.

In the AV Misaligned condition of the present study, the “spotlight” of top-down spatial attention had to be either divided or broadened across hemifields in order to successfully integrate the Target Talker’s face and voice. In vision, the spotlight of spatial attention can be efficiently divided into multiple locations across or within hemifields.^52–54^ Dividing auditory spatial attention, on the other hand, has been shown to come at a processing cost.^55^ Thus, it is possible that dividing cross-modal attention in the AV Misaligned condition decreased participants’ ability to track the Target Talker’s speech. However, in Exp. 2, a similar division of cross-modal attention was also required in the A-only Fixation Reversed condition; participants had to listen to speech in one hemifield while visually monitoring the other to detect the fixation catch trials. This did cause a performance decrement compared to the A-only Lateralized condition (in which participants listened to and visually monitored the same hemifield), but this effect was smaller than the difference between the AV Aligned and AV Misaligned conditions. Thus, dividing cross-modal spatial attention appeared to have an especially deleterious effect on the selective integration of AV speech. This is consistent with previous studies demonstrating that top-down attention is required for many forms of multisensory integration (see Talsma et al., 2010^46^ for review), as well as studies showing a reduction in behavioral^56^ and neural^57^ signatures of multisensory integration under a high degree of attentional load.

### When do we make use of auditory spatial information?

In auditory selective attention tasks, participants do not obligatorily rely on spatial features to separate component sounds from an auditory mixture. Top-down attention can be volitionally directed toward other sound features, such as pitch, depending on task demands and participant goals.^58^ Further, even when a target is defined only by a cued location, neural signatures of spatial attention appear at first, but are not sustained past initial selection of the relevant auditory target as long as the competing streams can be segregated on the basis of pitch.^59^ Both competing talkers were female in the present study, and so their voices were somewhat similar (although clearly differentiable) in fundamental frequency. Beyond F0 though, many other features were available to separate the two voices, including talker-specific characteristics (e.g. speech rate, vowel space, etc.), and in the AV conditions, the fact that each voice was temporally coherent with a separate face. Thus, cross-modal spatial features were by no means required to segregate the Target and Distractor Talkers, and yet we observed strong benefits of AV alignment nonetheless.

This discrepancy may trace its roots to the high degree of spatial reliability of the visual component of AV speech. The multisensory perceptual system is highly sensitive to the reliability of individual cues, such that vision dominates in instances of cross-modal spatial conflict (i.e. ventriloquism) as long as it remains more spatially reliable than audition.^60–62^ This visual dominance is distinct from auditory spatial information being ignored altogether. For instance, ventriloquism breaks down if the auditory and visual signals become too far separated in space, indicating that the auditory spatial information is still being encoded.^63^ Because part of the AV stimulus is in general spatially reliable, cross-modal spatial alignment may play a more obligatory role in AV than auditory-only selective attention, even if – as in the current study – spatial information is not explicitly task-relevant.

### Eye position effects on auditory spatial attention

The direction of eye gaze influences auditory responses in many stations along the processing hierarchy, including the inferior colliculus^64^, superior colliculus^65^, and primary auditory cortex.^66,67^ These effects are mirrored by improved auditory selective attention when fixating in the direction of an auditory target,^32,33^ which was also found in the A-only conditions of Exp. 2. The A-only Fixation Reversed condition created a particularly challenging scenario, as participants were asked to fix their gaze in the direction of the Distractor Talker. This has been shown to reduce the difference between target and distractor ERPs – a measure of successful deployment of attention – relative to fixation on the auditory target or a neutral location.^68^

Importantly, however, our ability to claim that participants were fixated on the exact location of the auditory source is limited because of the methodologies employed. First, sounds were spatialized with generalized HRTFs and presented through unknown headphones, so the degree to which participants externalized the sounds to the intended locations likely varied widely. Second, the use of HRTFs in the first place may have led to a weaker auditory spatial percept than free-field sound sources would have.^69^ Finally, without the use of eye-tracking we cannot be completely sure that participants maintained correct fixation throughout the experiment, although our fixation check task excluded several participants who likely failed to do so. Given these factors, the effects of eye position in this study may reflect the relationship between the hemifield of fixation and the hemifield of the Target Talker’s voice, more so effects of looking at the auditory target *per se*. Similarly, it is impossible to control factors such as display size or participant distance from the display in an online experiment format. Thus, effects of AV spatial alignment in this study should be interpreted at the hemifield level; further research using free-field sound sources, preferably in a laboratory setting so that eye gaze can be more reliably monitored, could distinguish the effects of integrating cross-modal information across hemifields from subtler forms of AV spatial misalignment.

## Conclusions

We demonstrated an improved ability to selectively attend AV speech when each talker’s face and voice were spatially aligned. Such an improvement was not found when corresponding faces and voices were spatially misaligned. This occurred despite the presence of alternative features, such as voice pitch and other talker-specific characteristics, that could be used to separate the competing speech streams. Taken together, these data provide evidence that cross-modal spatial alignment provides an important cue to the integration of AV speech stimuli in an acoustically and attentionally challenging environment.

## Acknowledgements

The authors would like to thank Dr. Tyler Perrachione for advice on mixed model construction and interpretation, and Audra Irvine for logistical support in collecting the online data. This work was supported by the Office of Naval Research (Grant N000141812069).

## Author Contributions

JTF, RKM, and BGSC all contributed to conceptualization and design of the experiments. Additionally, JTF implemented the experiments, collected, analyzed, and interpreted the data, and wrote the manuscript. RKM and BGSC also contributed to analysis and edited the manuscript.

## Additional Information (Competing Interests Statement)

The authors declare no competing interests.

## References

1. Erber, N. P. Auditory-visual perception of speech. J. Speech Hear. Disord. 40, 481–492 (1975).

2. Sumby, W. H. & Pollack, I. Visual contribution to speech intelligibility in noise. J. Acoust. Soc. Am. 26, 212–215 (1954).

3. Crosse, M. J., Di Liberto, G. M. & Lalor, E. C. Eye can hear clearly now: Inverse effectiveness in natural audiovisual speech processing relies on long-term crossmodal temporal integration. J. Neurosci. 36, 9888–9895 (2016).

4. Grant, K. W. & Seitz, P.-F. The use of visible speech cues for improving auditory detection of spoken sentences. J. Acoust. Soc. Am. 108, 1197–1208 (2000).

5. MacLeod, A. & Summerfield, Q. Quantifying the contribution of vision to speech perception in noise. Br. J. Audiol. 21, 131–141 (1987).

6. Ross, L. A., Saint-Amour, D., Leavitt, V. M., Javitt, D. C. & Foxe, J. J. Do you see what I am saying? Exploring visual enhancement of speech comprehension in noisy environments. Cereb. Cortex 17, 1147–1153 (2006).

7. Schwartz, J. L., Berthommier, F. & Savariaux, C. Seeing to hear better: Evidence for early audio-visual interactions in speech identification. Cognition 93, B69–B78 (2004).

8. Maddox, R. K., Atilgan, H., Bizley, J. K. & Lee, A. K. Auditory selective attention is enhanced by a task-irrelevant temporally coherent visual stimulus in human listeners. eLife; doi:10.7554/eLife.04995 (2015).

9. van Wassenhove, V., Grant, K. W. & Poeppel, D. Temporal window of integration in auditory-visual speech perception. Neuropsychologia; doi:10.1016/j.neuropsychologia.2006.01.001 (2007).

10. Besle, J., Fort, A., Delpuech, C. & Giard, M. H. Bimodal speech: Early suppressive visual effects in human auditory cortex. Eur. J. Neurosci.; doi:10.1111/j.1460-9568.2004.03670.x (2004).

11. van Wassenhove, V., Grant, K. W. & Poeppel, D. Visual speech speeds up the neural processing of auditory speech. Proc. Natl. Acad. Sci.; doi:10.1073/pnas.0408949102 (2005).

12. Peelle, J. E. & Sommers, M. S. Prediction and constraint in audiovisual speech perception. Cortex 68, 169–181 (2015).

13. Simon, D. M. & Wallace, M. T. Integration and Temporal Processing of Asynchronous Audiovisual Speech. J. Cogn. Neurosci. 30, 319–337 (2018).

14. Bertelson, P., Vroomen, J., Wiegeraad, G. & de Gelder, B. Exploring the relation between McGurk interference and ventriloquism. Proc. ICSLP 559–562 (1994).

15. Jones, J. A. & Munhall, K. G. Effects of separating auditory and visual sources on audiovisual integration of speech. Can. Acoust. 25, 13–19 (1997).

16. DeLoss, D. J. & Andersen, G. J. Aging, spatial disparity, and the sound-induced flash illusion. PLoS ONE 10, e0143773 (2015).

17. Innes-Brown, H. & Crewther, D. The impact of spatial incongruence on an auditory-visual illusion. PLoS ONE 4, e6450 (2009).

18. Bizley, J. K., Shinn-Cunningham, B. G. & Lee, A. K. C. Nothing is irrelevant in a noisy world: Sensory illusions reveal obligatory within-and across-modality integration. J. Neurosci. 32, 13402–13410 (2012).

19. Van der Burg, E., Olivers, C. N. L., Bronkhorst, A. W. & Theeuwes, J. Pip and pop: Nonspatial auditory signals improve spatial visual search. J. Exp. Psychol. Hum. Percept. Perform. 34, 1053–1065 (2008).

20. Fleming, J. T., Noyce, A. L. & Shinn-Cunningham, B. G. Audio-visual spatial alignment improves integration in the presence of a competing audio-visual stimulus. Neuropsychologia 146; doi:10.1016/j.neuropsychologia.2020.107530 (2020).

21. Atilgan, H. et al. Integration of visual information in auditory cortex promotes auditory scene analysis through multisensory binding. Neuron 1–16 (2018).

22. Zion Golumbic, E., Cogan, G. B., Schroeder, C. E. & Poeppel, D. Visual input enhances selective speech envelope tracking in auditory cortex at a “cocktail party”. J. Neurosci. 33, 1417–1426 (2013).

23. Litovsky, R. Y. Spatial release from masking. Acoust. Today 8, 18–25 (2012).

24. Watson, C. S. Uncertainty, informational masking, and the capacity of immediate auditory memory. in Auditory processing of complex sounds 267–277 (1987).

25. Wu, X. et al. The effect of perceived spatial separation on informational masking of Chinese speech. Hear. Res. 199, 1–10 (2005).

26. Durlach, N. I. et al. Note on informational masking (L). J. Acoust. Soc. Am. 113, 2984 (2003).

27. Helfer, K. S. & Freyman, R. L. The role of visual speech cues in reducing energetic and informational masking. J. Acoust. Soc. Am. 117, 842–849 (2005).

28. Chait, M., Poeppel, D. & Simon, J. Z. Neural Response Correlates of Detection of Monaurally and Binaurally Created Pitches in Humans. Cereb. Cortex 16, 835–848 (2006).

29. Cramer, E. M. & Huggins, W. H. Creation of Pitch through Binaural Interaction. J. Acoust. Soc. Am. 30, 413–417 (1958).

30. Milne, A. E. et al. An online headphone screening test based on dichotic pitch. Behav. Res. Methods; doi:10.3758/s13428-020-01514-0 (2020).

31. Algazi, V. R., Duda, R. O., Thompson, D. M. & Avendaño, C. The CIPIC HRTF database. in Proceedings of the 2001 IEEE Workshop on the Applications of Signal Processing to Audio and Acoustics, Cat. No.01TH8575 99–102 (2001).

32. Maddox, R. K., Pospisil, D. A., Stecker, G. C. & Lee, A. K. C. Directing eye gaze enhances auditory spatial cue discrimination. Curr. Biol. 24, 748–752 (2014).

33. Reisberg, D., Scheiber, R. & Potemken, L. Eye position and the control of auditory attention. J. Exp. Psychol. Hum. Percept. Perform. 7, 318–323 (1981).

34. Cui, Q. N., Razavi, B., O’Neill, W. E. & Paige, G. D. Perception of auditory, visual, and egocentric spatial alignment adapts differently to changes in eye position. J. Neurophysiol. 103, 1020–1035 (2010).

35. Razavi, B., O’Neill, W. E. & Paige, G. D. Auditory spatial perception dynamically realigns with changing eye position. J. Neurosci. 27, 10249–10258 (2007).

36. Senkowski, D., Saint-Amour, D., Höfle, M. & Foxe, J. J. Multisensory interactions in early evoked brain activity follow the principle of inverse effectiveness. NeuroImage 56, 2200–2208 (2011).

37. Stevenson, R. A. et al. Inverse effectiveness and multisensory interactions in visual event-related potentials with audiovisual speech. Brain Topogr. 25, 308–326 (2012).

38. Stevenson, R. A. & James, T. W. Audiovisual integration in human superior temporal sulcus: Inverse effectiveness and the neural processing of speech and object recognition. NeuroImage 44, 1210–1223 (2009).

39. Altieri, N. A., Pisoni, D. B. & Townsend, J. T. Some normative data on lip-reading skills (L). J. Acoust. Soc. Am. 130, 1–4 (2011).

40. Kidd, G., Mason, C. R., Rohtla, T. L. & Deliwala, P. S. Release from masking due to spatial separation of sources in the identification of nonspeech auditory patterns. J. Acoust. Soc. Am. 104, 422–31 (1998).

41. Marrone, N., Mason, C. R. & Kidd, G. The effects of hearing loss and age on the benefit of spatial separation between multiple talkers in reverberant rooms. J. Acoust. Soc. Am. 124, 3064–3075 (2008).

42. Shinn-Cunningham, B. G., Ihlefeld, A., Satyavarta & Larson, E. Bottom-up and top-down influences on spatial unmasking. Acta Acust. United Acust. 91, 967–979 (2005).

43. Colonius, H. & Diederich, A. The race model inequality: interpreting a geometric measure of the amount of violation. Psychol. Rev. 113, 148–154 (2006).

44. Fraser, S., Gagné, J.-P., Alepins, M. & Dubois, P. Evaluating the effort expended to understand speech in noise using a dual-task paradigm: The effects of providing visual speech cues. J. Speech Lang. Hear. Res. 53, 18–33 (2010).

45. Strand, J. F., Brown, V. A. & Barbour, D. L. Talking points: A modulating circle increases listening effort without improving speech recognition in young adults. Psychon. Bull. Rev. 27, 536–543 (2020).

46. Talsma, D., Senkowski, D., Soto-Faraco, S. & Woldorff, M. G. The multifaceted interplay between attention and multisensory integration. Trends Cogn. Sci. 14, 400–410 (2010).

47. McGurk, H. & Macdonald, J. Hearing lips and seeing voices. Nature 264, 746–748 (1976).

48. Andersen, T. S., Tiippana, K., Laarni, J., Kojo, I. & Sams, M. The role of visual spatial attention in audiovisual speech perception. Speech Commun. 51, 184–193 (2009).

49. Tiippana, K., Puharinen, H., Möttönen, R. & Sams, M. Sound location can influence audiovisual speech perception when spatial attention is manipulated. Seeing Perceiving 24, 67–90 (2011).

50. Talsma, D. & Woldorff, M. G. Selective attention and multisensory integration: multiple phases of effects on the evoked brain activity. J. Cogn. Neurosci. 17, 1098–1114 (2005).

51. Fairhall, S. L. & MacAluso, E. Spatial attention can modulate audiovisual integration at multiple cortical and subcortical sites. Eur. J. Neurosci. (2009) doi:10.1111/j.1460-9568.2009.06688.x.

52. Malinowski, P., Fuchs, S. & Müller, M. M. Sustained division of spatial attention to multiple locations within one hemifield. Neurosci. Lett. 414, 65–70 (2007).

53. McMains, S. A. & Somers, D. C. Processing efficiency of divided spatial attention mechanisms in human visual cortex. J. Neurosci. 25, 9444–9448 (2005).

54. Müller, M. M., Malinowski, P., Gruber, T. & Hillyard, S. A. Sustained division of the attentional spotlight. Nature 424, 309–312 (2003).

55. Parasuraman, R. Auditory evoked potentials and divided attention. Psychophysiology 15, 460–465 (1978).

56. Alsius, A., Navarra, J., Campbell, R. & Soto-Faraco, S. Audiovisual integration of speech falters under high attention demands. Curr. Biol. 15, 839–843 (2005).

57. Alsius, A., Möttönen, R., Sams, M. E., Soto-Faraco, S. & Tiippana, K. Effect of attentional load on audiovisual speech perception: evidence from ERPs. Front. Psychol. 5, (2014).

58. Maddox, R. K. & Shinn-Cunningham, B. G. Influence of task-relevant and task-irrelevant feature continuity on selective auditory attention. J. Assoc. Res. Otolaryngol. 13, 119–129 (2012).

59. Bonacci, L. M., Bressler, S. & Shinn-Cunningham, B. G. Nonspatial features reduce the reliance on sustained spatial auditory attention. Ear Hear. 41, 1635–1647 (2020).

60. Recanzone, G. H. Rapidly induced auditory plasticity: The ventriloquism aftereffect. Proc. Natl. Acad. Sci. 95, 869–875 (1998).

61. Wozny, D. R. & Shams, L. Recalibration of auditory space following milliseconds of cross-modal discrepancy. J. Neurosci. Off. J. Soc. Neurosci. 31, 4607–12 (2011).

62. Alais, D. & Burr, D. The ventriloquist effect results from near-optimal bimodal integration. Curr. Biol. 14, 257–262 (2004).

63. Bosen, A. K. et al. Comparison of congruence judgment and auditory localization tasks for assessing the spatial limits of visual capture. Biol. Cybern. (2016) doi:10.1007/s00422-016-0706-6.

64. Groh, J. M., Trause, A. S., Underhill, A. M., Clark, K. R. & Inati, S. Eye position influences auditory responses in primate inferior colliculus. Neuron 29, 509–518 (2001).

65. Jay, M. F. & Sparks, D. L. Auditory receptive fields in primate superior colliculus shift with changes in eye position. Nature 309, 345–347 (1984).

66. Fu, K.-M. G. et al. Timing and laminar profile of eye-position effects on auditory responses in primate auditory cortex. J. Neurophysiol. 92, 3522–3531 (2004).

67. Werner-Reiss, U., Kelly, K. A., Trause, A. S., Underhill, A. M. & Groh, J. M. Eye position affects activity in primary auditory cortex of primates. Curr. Biol. CB 13, 554–562 (2003).

68. Okita, T. & Wei, J.-H. Effects of eye position on event-related potentials during auditory selective attention. Psychophysiology 30, 359–365 (1993).

69. Brungart, D. S. & Simpson, B. D. Auditory localization of nearby sources in a virtual audio display. in Proceedings of the 2001 IEEE Workshop on the Applications of Signal Processing to Audio and Acoustics (Cat. No.01TH8575) 107–110 (IEEE, 2001). doi:10.1109/ASPAA.2001.969554.

